# Development of a Real-Time Neural Controller using an EMG-Driven Musculoskeletal Model

**DOI:** 10.1101/2024.12.06.627232

**Authors:** Joel Biju Thomas, Brokoslaw Laschowski

## Abstract

Here we present our development of a novel real-time neural controller based on an EMG-driven musculoskeletal model, designed for volitional control of robots and computers. Our controller uniquely enables motion control during both isometric and non-isometric muscle contractions. We address several key challenges in EMG control system design, including accuracy, latency, and robustness. Our approach combines EMG signal processing, neural activation dynamics, and Hill-type muscle modeling to translate neural commands into muscle forces, which can enhance robustness against electrode variability and signal noise. Additionally, we integrate muscle activation dynamics with impedance control, inspired by the human motor control system, for smooth and adaptive interactions. As an initial proof of concept, we demonstrated that our system could control a robot actuator across a range of movements, both static and dynamic, and at different operating speeds, achieving high reference tracking performance and state-of-the-art processing times of 2.9 ms, important for real-time embedded computing. This research helps lay the groundwork for next-generation neural-machine interfaces that are fast, accurate, and adaptable to diverse users and control applications.

## I. Introduction

Surface electromyography (EMG) is a non-invasive neural interface that can be used to volitionally control robots and computers by translating feedforward neural commands from the brain (i.e., that encode information about motor intent) into actionable control signals. Despite this potential, the development of reliable EMG neural controllers faces significant challenges, particularly in terms of system accuracy, latency, and adaptability to diverse users [1]. Applications in patients with amputation require precise control during isometric muscle contractions, while controllers for able-bodied persons need to also support dynamic, non-isometric contractions for natural, functional movements. High accuracy and low latency, which are critical to optimal user experience, remains an open problem for these neural control systems [2]-[5].

Two of the main paradigms used for EMG neural control are data-driven classification models and neuromusculoskeletal models. Data-driven methods typically use machine learning algorithms to classify, or decode, the joint movement from EMG signals [6]-[7] such as quadratic discriminant analysis used by [6] for patients with amputation. While these models have shown success, they often make simplifying assumptions about the underlying data distributions, which can limit their robustness given the inherent non-Gaussian nature of EMG data. Factors such as noise, muscle fatigue, and inter-subject variability also pose significant challenges to these classification models for long-term operation [7]-[8], including requiring long user training times. More complex models such as convolutional or recurrent neural networks could help to eliminate these assumptions, but with the potential trade-off of comprising real-time inference with low latency.

In contrast, neuromusculoskeletal modelling approaches model muscle forces and joint dynamics more directly, thus allowing for improved accuracy in muscle force prediction and system adaptability. Previous research [5], [8] has used these computational models to simulate muscle-generated forces from EMG signals. However, they tend to lack an EMG-to-activation model, which can limit their robustness to variations in sensor placement and EMG signal-to-force mapping accuracy, constraining their practical utility. Despite progress in both machine learning and neuromusculoskeletal modelling EMG controllers, the focus has been on isometric control for amputees, with less attention given to the general population (i.e., able-bodied users) and to dynamic, non-isometric movements [5]-[7], which can be more technically challenging.

To address these gaps, here we present our development of a novel EMG neural controller driven by a musculoskeletal model. We include an EMG-to-activation model to improve robustness and to more accurately map EMG signals to muscle forces. Our controller uniquely handles both isometric and non-isometric muscle contractions, expanding functionality over previous work. By combining the strengths of neuromusculoskeletal modeling with a more accurate EMG signal-to-force mapping model, our research aims to lay the foundation for next-generation neural-machine interfaces that can support diverse users and control applications. While the focus of this study was on the engineering control system design, as an initial proof-of-concept, we demonstrate that our EMG neural controller can be used to accurately and in real-time control a robotic actuator across a range of operating speeds using neural commands to leg muscles (see Fig. 1).

**Fig. 1.**
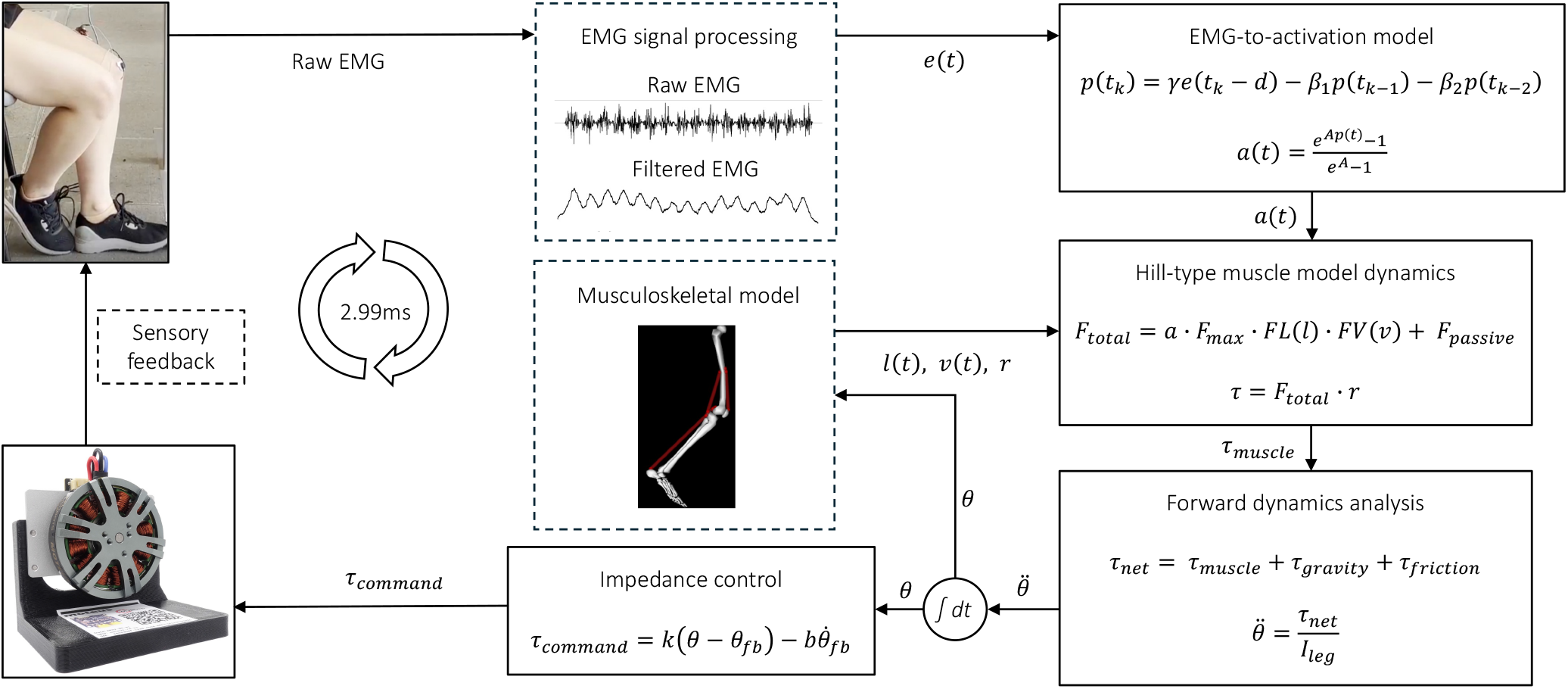
System diagram of our real-time neural controller using an EMG-driven musculoskeletal model. We process and convert raw EMG signals into neural activations, which drive our muscle and musculoskeletal dynamics models to produce joint torques and movement. We also developed an impedance controller to generate torque commands for our motor actuator, which takes as input the simulated joint kinematics.

## II. Methods

### A. Signal Processing

We used standard EMG processing to enhance our signal quality and to reduce unwanted artifacts [9], including a 200x gain amplifier, bandpass filter between 20-498 Hz, signal rectification, and offset removal. We also performed envelope detection with a frequency response centered at 3.6 Hz, capturing the slow-varying envelope of our rectified EMG signal [9]. To further enhance clarity, we amplified the envelope using a secondary gain amplifier and digitized the signal with an Arduino Uno R3 as an ADC, sampling at 1400 Hz. The digitized EMG signals were sent to Raspberry PI using USB UART for further processing. We normalized the signal to a 0*–*1 range to enable consistency across subjects using maximal voluntary contraction (MVC) and minimum EMG amplitude during full muscle flexion and relaxation. This minimizes variations in amplitude due to individual physiology, which allow for more accurate cross-subject and experimental comparisons [10].

### B. Neural Activation Model

The biological tissue between the EMG electrode and muscle naturally filters the neural signals produced by the muscles before they are recorded. This can pose an issue where factors such as the positioning of the electrodes, ambient temperature, skin condition, and electrical impedance can influence filtering [11]. To reduce this effect of tissue filtering, we developed a new EMG-to-activation model to interpret the EMG signals and estimate muscle force, accounting for physiological delays and nonlinearities in muscle activations, enhancing precision and reliability.

Our new EMG-to-activation model begins with activation dynamics, which transforms the processed EMG signals e(t) into neural activations p(t) [12]. This process addresses the time delay between EMG onset and muscle force generation, as well as the longer durations of muscle forces compared to EMG signals [11]. Unlike previous work [5], our controller includes this EMG-to-activation model for a more reliable muscle force estimation by taking into account biological delays and nonlinearities. We used a critically-damped linear second-order differential system to model the delayed muscle twitch response, which occurs when a single action potential stimulates an individual muscle fiber [12]. In its discrete form, see Equation 1, our second-order system can be expressed recursively to compute p(t), providing a more accurate representation of the muscle’s force generation based on the input EMG neural signal.

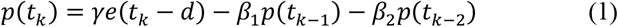

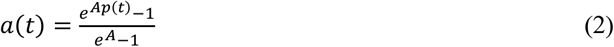

where *d* is the electromechanical delay and γ, *β*_1_, and *β*_2_ are coefficients governing the second-order system dynamics. To ensure a positive and stable outcome, we enforced specific constraints: *β*_1_ *= C*_1_ + *C*_1_ and *β*2 *= C*_1_*C*_2_, with the conditions |*C*_1_| *<* 1 and |*C*_2_| *<* 1. Furthermore, to preserve the filter’s gain, we must satisfy γ *– β*_1_ *– β*_2_ *=* 1. We used a nonlinear conversion between the neural activations *p*(*t*) and muscle activations *a*(*t*), see Equation 2, in order to address the nonlinear relationship between the EMG signal and muscle force generation at low-force and high twitch frequencies [12].

### C. Muscle Model

We developed a Hill-type muscle model to mathematically model the muscle force generation based on the relationships between muscle length, contraction speed, and activation dynamics. Our Hill-type muscle model combines active forces from neural activations with the passive forces from muscle stretching for a more biofidelic representation of biological muscle dynamics. We used the force-length factor to model how the muscle force generation depends on its length relative to optimal length, with maximum force at the optimal length and an exponential decrease as the muscle length deviates

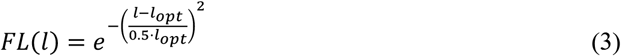

where FL(l) is the muscle force-length factor, l is the current muscle length, and *l*_*opt*_ is the optimal muscle length [13]. For concentric muscle contractions, where velocity is greater than or equal to 0, we used a linear force-velocity relationship. For eccentric muscle contractions, where the velocity is less than zero, we used a nonlinear force-velocity relationship.

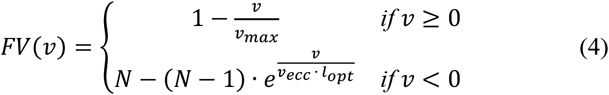

where *FV*(*v*) is the muscle force-velocity factor, *v* is the muscle velocity, *v*_*max*_ is the maximum shortening velocity, *N* is the eccentric muscle force factor, and *V*_*ecc*_ is the characteristic velocity of eccentric muscle contractions [13]. Active force *(F*_*active*_) is defined as the product of the muscle activation, maximum isometric force, force-length factor, and the force-velocity factor, and is represented by:

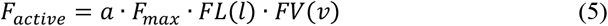

where *a* is the muscle activation level, ranging from 0 to 1, and *F*_*max*_ is the maximum isometric force. We derived muscle length and velocity from the joint angle and angular velocity using our musculoskeletal model. We calculated the muscle length by converting the joint angle into radians, adjusting for deviations from the reference angle, and scaling by moment arm. We calculated muscle velocity by converting the joint angular velocity into radians and scaling using the moment arm. Passive forces *(F*_*passive*_) are generated when the muscle stretches beyond its optimal length, increasing exponentially with stretch. We assumed no passive muscle force is produced below the optimal length, but beyond that, the force grows exponentially and is defined by the maximum isometric force and a passive stiffness coefficient *(C*_*passive*_) [13].

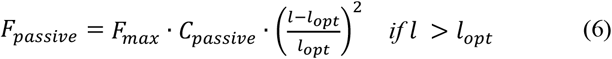

We calculated the total force *F*_*total*_ generated by the muscle using the sum of the active *F*_*active*_ and passive *F*_*passive*_ muscle forces. The resultant joint torque *τ* generated by the muscle was calculated using the product of the total force *F*_*total*_ and the moment arm *r*.

### D. Forward Dynamics

We used forward dynamics analysis to calculate the joint motion from the applied muscle torques, gravitational forces, and joint friction in our musculoskeletal system model.

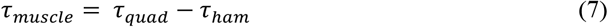

where *τ*_*muscle*_ is the net joint torque, which is the difference between the joint torques generated by the quadricep muscles *τ*_*quad*_ and the hamstring muscles *τ*_*ham*._ We estimated the joint torque generated from gravity acting on the leg *τ*_*gravity*_ using Equation 8:

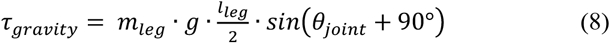

where *m*_*leg*_ is the estimated mass of the leg, *g* is acceleration due to gravity, *l*_*leg*_ is the length of the leg, and *θ*_*joint*_ is the instantaneous joint angle. We estimated the frictional torque in the joint using *τ*_*friction*_ *= −*b,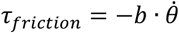 where *b* is the biological damping coefficient and 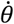 is the angular velocity of the joint. The net joint torque *τ*_*net*_ is the sum of *τ*_*muscle*_, *τ*_*gravity*_ and *τ*_*friction*_, and the resulting angular acceleration 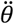 is calculated as 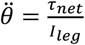 where *l*_*leg*_ is the estimated moment of inertia of the leg. When calculating *τ*_*net*_ for isometric contractions, we set *τ*_*gravity*_ and the muscle velocity to 0. We did this so that the EMG signal is used to generate an angular velocity, which means that the motor will only actuate when the muscles are contracting and is stationary when the muscles are relaxed. This is important for downstream applications in patients with amputation, for example, to minimize fatigue while operating our controller. We assume a constant muscle length.

### E. Kinematics

We used kinematic analysis to continuously track the joint angular velocity and position over time based on the initial conditions and integrals. We calculated joint angular velocity 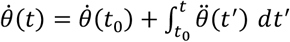 using the initial velocity 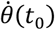 and the cumulative angular acceleration 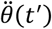. We calculated the joint angular position 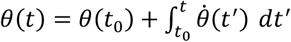 using the initial position *θ*(*t*_o_) and the integrated velocity.

### F. Impedance Control

We developed an impedance controller for our lower-level controller to convert joint motion (input) to torque (output). Impedance control is widely used in robotics to control the interaction between force and motion by adjusting mechanical impedance. This control paradigm is inspired by human motor control, where the nervous system regulates muscle stiffness and compliance for smooth and adaptive physical interactions with the environment. Research has consistently shown that myoelectric impedance control outperforms conventional velocity-based control in constrained motion tasks [4]. Our impedance controller is expressed as:

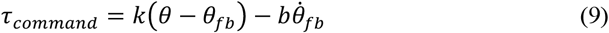

where *τ*_command_ is the torque command to the field-oriented controller of our motor actuator, *θ* is the joint position of our musculoskeletal model, *θ*_fb_ is the motor position, 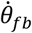 is motor velocity, and *k* and *b* are stiffness and damping, respectively. Here we assumed that *k* and *b* are fixed constants that do not change based on muscular co-contraction.

### G. Experiments

As a proof of concept to evaluate the performance of our new EMG neural controller, we tested the ability of a user to volitionally control a robotic actuator, specifically a brushless DC electric motor, during isometric (static) and non-isometric (dynamic) muscle contractions. We determined the accuracy of our controller based on the root mean square error (RMSE) with reference joint angle trajectories, and we measured latency to assess responsiveness and real-time control.

We prepared the EMG recording sites similar to [2]. To locate the best electrode position, we had the user fully extend their leg and perform maximum quadricep contractions while we palpated the muscle. Two electrodes, ∼25 mm apart, were placed on the quadricep muscle. We used a similar procedure for the hamstrings when the knee was flexed. We had the user perform maximum voluntary contraction and relaxation of their quadricep and hamstring muscles and we recorded the maximum and minimum EMG signals for normalization. In a seated posture, we instructed the user to flex and extent their instrumented leg with the goal of controlling the motor angle to follow a reference video displayed on a computer. The user practiced with our controller for ∼10 minutes before results were recorded. The user performed non-isometric, sinusoidal knee flexion and extension, followed by isometric contractions with a fixed knee joint.

We evaluated the accuracy of our EMG neural controller by calculating the RMSE between the neural-controlled motor angles as measured by our motor controller and the calculated knee joint angles from the reference video displayed on the computer. We conducted 80-second trials to test different movement frequencies at 0.2 Hz, 0.25 Hz, 0.33 Hz, and 0.5 Hz, with RMSE selected for direct comparisons with the previous state-of-the-art [6]. We developed the reference video by recording the user flex and extent their leg at the specified frequencies, guided by a metronome. We calculated the joint angles using Vicon motion tracker with 16 passive markers on the leg sampled at 100 Hz and the Vicon ProCalc software. Lastly, we measured our controller’s latency by combining the processing times from the Arduino and Raspberry Pi during a motor controller update.

## III. Results

Table 1 shows the accuracy of our EMG neural controller across the different movement frequencies, ranging from 0.50 Hz (2 seconds) to 0.20 Hz (5 seconds). For isometric muscle contractions, our controller achieved the lowest RMSE of 8.2° at 0.25 Hz and the highest RMSE of 9.7° at 0.33 Hz (Fig. 2). Non-isometric muscle contractions followed a similar trend, with the lowest RMSE of 7.9° at 0.25 Hz and highest RMSE of 9.7° at both 0.33 Hz and 0.20 Hz (Fig. 3). Our controller achieved an average RMSE of 9.2° across all conditions, demonstrating a relatively consistent accuracy for a diverse set of motion control. Moreover, our controller achieved a state-of-the-art end-to-end processing time of 2.9 ms, showcasing its responsiveness and real-time control.

**Table 1.**
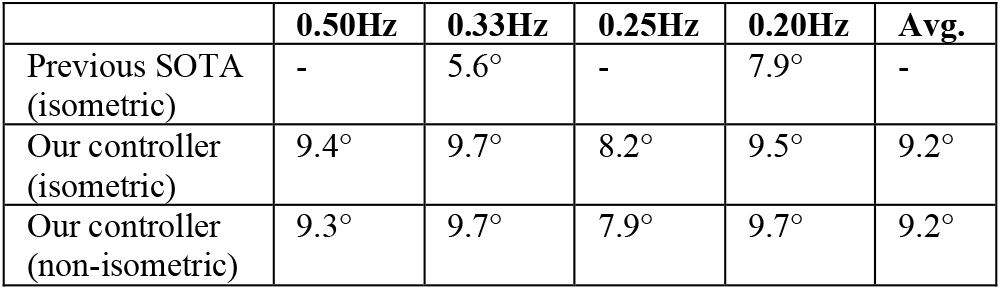
Comparison of our EMG neural controller with the previous state-of-the-art [6]. We evaluated performance based on the RMSE in joint angles for isometric and non-isometric muscle contractions across different movement frequencies.

|  | 0.50Hz | 0.33Hz | 0.25Hz | 0.20Hz | Avg. |
| --- | --- | --- | --- | --- | --- |
| Previous SOTA (isometric) | - | 5.6° | - | 7.9° | - |
| Our controller (isometric) | 9.4° | 9.7° | 8.2° | 9.5° | 9.2° |
| Our controller (non-isometric) | 9.3° | 9.7° | 7.9° | 9.7° | 9.2° |

**Fig. 2.**
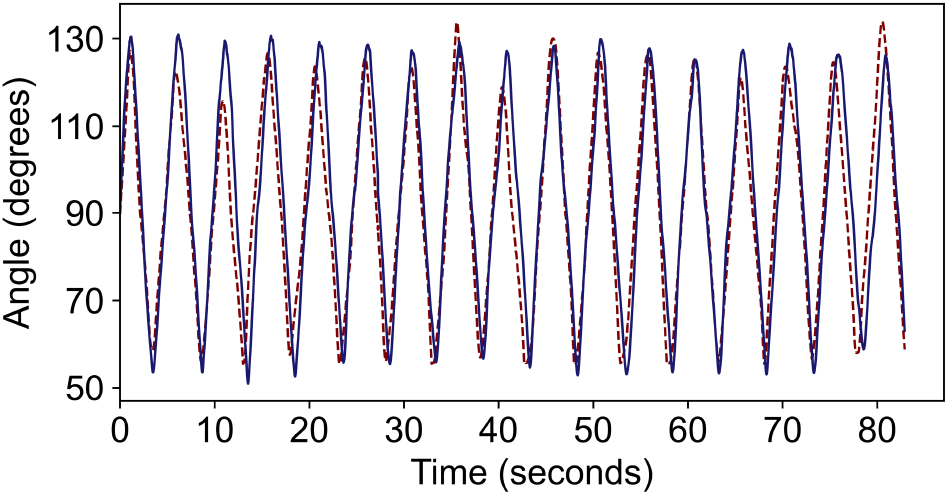
Trajectory tracking of our EMG neural controller for isometric, sinusoidal movements at 0.2 Hz frequency. The red dashed signal is the reference joint angle, and the blue dashed signal is the motor angle of our neural-controlled actuator.

**Fig. 3.**
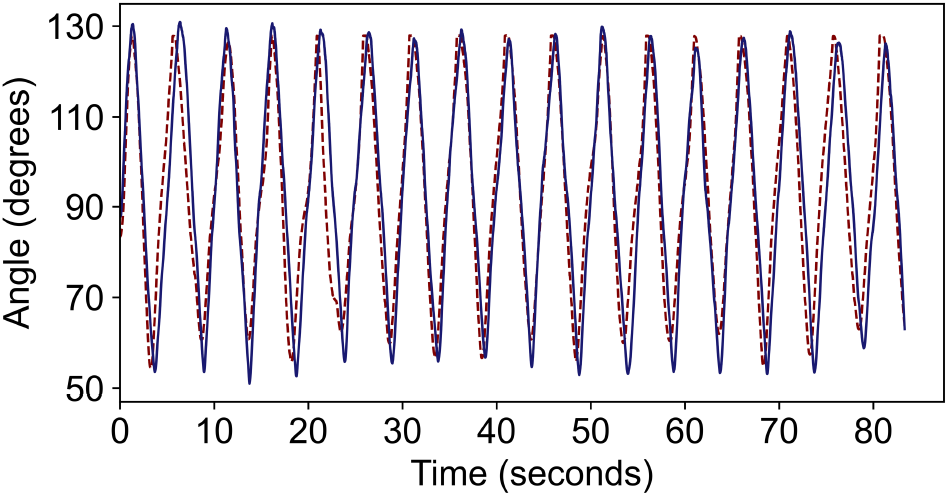
Trajectory tracking of our EMG-driven neural controller for nonisometric, sinusoidal movements at 0.2 Hz frequency. The red dashed signal is the reference joint angle, and the blue dashed signal is the motor angle of our neural-controlled actuator.

## IV. Discussion

In this study, we developed a new real-time neural controller using an EMG-driven musculoskeletal model. Our system enables intuitive and responsive control for both isometric and non-isometric movements. Using processed EMG signals, we dynamically drive a muscle model to translate the user’s intent into precise motor control with minimal latency. As an initial proof-of-concept, we demonstrated that our system could use feedforward neural commands to the leg muscles for accurate real-time control of a robot actuator across a range of different movements and operating speeds. This is partially due to our incorporation of muscle activation dynamics and a Hill-type muscle model with impedance control. Motivated by these promising results, our controller could be used to support the development of next-generation neural-machine interfaces for diverse users and control applications.

Our EMG neural controller achieved an average RMSE of 9.2° over 80-second experiments for both isometric and non-isometric movements. In comparison, [6] achieved 6.2° RMSE in patients with leg amputation. For 3-and 5-second isometric movement frequencies, our controller achieved an RMSE of 9.7° and 9.5°, whereas [6] achieved an RMSE of 5.6° and 7.9°, respectively. Although we achieved slightly lower accuracy, our controller can uniquely support both isometric and non-isometric motion control, whereas [6] focused on patients with amputation, and thus limited to isometric movements. Given that joint proprioception accuracy in healthy adults is ∼2.5° [14], our controller’s instantaneous error difference is arguably imperceptible, especially as RMSE reflects cumulative error. Our study is one of the few to develop an EMG neural controller for both isometric and non-isometric movements. Moreover, our use of a neuromusculoskeletal model has the ability to enhance our system’s robustness compared to [6], which used a data-driven model. Our controller also requires significantly less training time, achieving intuitive control with only 10 minutes of user familiarization, whereas [6] required one-hour of training per session, repeated three times a week. Most notably, our controller achieved state-of-the-art processing speeds of 2.9 ms, compared to 10 ms times for other two-channel EMG systems [3].

Despite these advantages, our controller has limitations. It cannot not seamlessly transition between isometric and non-isometric states as it lacks a learning algorithm for movement classification, operating in discrete controllers rather than continuously. Moreover, while our neuromusculoskeletal model offers flexibility via tunable parameters, this can lead to inconsistent accuracy across users such that optimizing our model for one user may yield suboptimal results for another unless calibrated. While muscle co-contractions are common in patients with amputation [6], our controller does not include signal processing to account for this, which could otherwise benefit our controller. Although our EMG-to-activation model can theoretically improve system robustness to uncertainty in electrode placement, we did not explicitly test this. Another limitation of our study was that the experiments were based on one user. However, the focus of this research was on the engineering control system design, while the user testing was more to demonstrate an initial proof-of-concept.

Future research could expand our control system design to also include a classification algorithm to detect isometric and non-isometric movements, advancing toward a more universal EMG neural controller. Incorporating additional processing to manage muscle co-contraction—such as principal component analysis used by [6], or unsupervised learning algorithms like k-means clustering—could improve accuracy. Other neural sensors such as electroencephalography (EEG) could also be used to decode different movements and tasks [15]-[16]. Adding sensors like accelerometers or computer vision could help supplement our neuromusculoskeletal model by providing data on gravitational directionality for better control in variable orientations [17]-[21]. Overall, future research is needed to explore generalizability of our control system to novel users and tasks.

In conclusion, we developed a new EMG neural controller driven by neuromusculoskeletal model. As a proof-of-concept, we showed that our controller could achieve accurate and consistent motion control of a robot actuator across a range of different movements and operating speeds, in addition to state-of-the-art processing speeds, which is important for real-time embedded computing. With further development, our controller could be used to interact with and control robots and computers, providing an intuitive neural-machine interface.

## Acknowledgment

This research study is dedicated to the people of Ukraine in response to Russia’s ongoing invasion. We thank Stephen Yang and Yume Yamamoto for their assistance.

